# Age Differences in Hemispheric Lateralization in Spatial and Verbal Visual Working Memory

**DOI:** 10.1101/577858

**Authors:** Myriam C. Sander, Patrizia M. Maier, Natan Napiórkowski, Kathrin Finke, Thomas Töllner, Hermann J. Müller, Ulman Lindenberger, Markus Werkle-Bergner, Iris Wiegand

**Author notes:** Correspondence should be addressed to Myriam Sander or Iris Wiegand.

## Abstract

Due to hemispheric specialization of the human brain, neural signatures of visual working memory (WM) performance are expected to differ between tasks involving verbal versus spatial memoranda. Theories of cognitive aging suggest a reduction of hemispheric specialization in older adults. Using behavioral and neural WM capacity markers, we assessed hemispheric lateralization in younger and older adults performing a spatial or verbal visual WM task. Participants encoded information presented in the left or right hemifield. We observed behavioral advantages for spatial stimuli processed in the right hemisphere and for verbal stimuli processed in the left hemisphere. While younger adults showed lateralization in both tasks, older adults showed lateralization only in the verbal task. Lateralization was assessed by the contralateral delay activity (CDA) on the neural level. CDA amplitudes displayed hemispheric lateralization for verbal versus spatial material, but this effect was age-invariant. While our findings support right-hemispheric specialization for spatial information maintenance, and left-hemispheric specialization for verbal information maintenance, we could not confirm a generalized reduction in hemispheric lateralization at older ages.

## Introduction

Working memory (WM) is our ability to encode information from our surroundings, maintain it over a short period of time, and use it for future behavior and task solving (e.g., Eriksson, Vogel, Lansner, Bergstrom, & Nyberg, 2015). On the neural level, WM is supported by a broad network of brain regions, encompassing parietal and frontal as well as subcortical areas (Erikson et al., 2015). Besides a core WM network, additional brain regions are recruited dependent on the type of to-be-represented information. For example, there is converging evidence that the left hemisphere is preferentially recruited for language-related functions, whereas the right hemisphere is strongly involved in visuospatial processing (e.g., Broca, 1861; Mesulam, 1981; Pujol et al., 1999; Smith & Jonides, 1998; Springer & Deutsch, 1998).

Neuropsychological studies showed that patients with lesions in the left hemisphere suffer more often from language disorders such as aphasia, whereas patients with lesions in the right hemisphere experience visuospatial attention deficits such as neglect (Corbetta & Shulman, 2011; Mesulam, 1981; Smith & Jonides, 1998). The right-hemispheric dominance for visuospatial information is further supported by a phenomenon termed ‘pseudoneglect’, a small but systematic attentional bias to the left hemifield shown by healthy individuals in visuospatial tasks (Benwell, Thut, Grant, & Harvey, 2014; Bowers & Heilmann, 1980; Finke et al., 2005; Newman, Loughnane, Kelly, O’Connell, & Bellgrove, 2017). Furthermore, neuroimaging studies found material-specific hemispheric lateralization in perceptual as well as WM tasks in healthy adults and patients (Cai, Van der Haegen, & Brysbaert, 2013; Koenigs, Barbey, Postle, & Grafman, 2009; Nee et al., 2013; Owen, McMillan, Laird, & Bullmore, 2005; Reuter-Lorenz et al., 2000; Shallice, 1988; Smith & Jonides, 1998, 1999; Thomason et al., 2008;). Specifically, WM tasks with verbal material activate primarily left-hemispheric areas in the posterior parietal cortex as well as Broca, premotor, and supplementary motor areas, while WM tasks with spatial material activate right-hemispheric posterior parietal, occipital, and frontal cortical areas. For a review of functional magnetic resonance imaging (fMRI) and positron emission tomography (PET) evidence, see Smith and Jonides (1998).

Senescent neural changes, such as reductions in gray and white matter in parietal and frontal brain regions, affect WM performance in older adults (Nyberg et al., 2014; see also Sander et al., 2012, for a review). However, the degree of age-related decline in WM performance may depend on the type of the to-be-maintained material. In particular, some studies reported stronger age-related decline in spatial compared to verbal WM (Jenkins, Myerson, Joerding, & Hale, 2000; Myerson, Emery, White, & Hale, 2003; but see Salthouse, 1995, and Park et al., 2002, for contradicting evidence) and other tasks (e.g., results from the Wechsler Adult Intelligence Scale, see Goldstein & Shelly, 1981). One account for these findings is the so-called ‘right hemi-aging model’ (RHAM) by Brown and Jaffe (1975). It proposes that the right hemisphere and associated functions show a faster age-related decline compared to the left hemisphere (Dolcos, Rice, & Cabeza, 2002; Goldstein & Shelly, 1981; Klisz, 1978). Accordingly, visuo-spatial tasks with a stronger reliance on the right hemisphere should be relatively more affected by aging than verbal tasks, which depend on left-hemispheric processing. An alternative account to explain age-related changes in lateralization is provided by the hemispheric asymmetry reduction in older adults (HAROLD) model by Cabeza (2002). The HAROLD model suggests generally reduced hemispheric differences in older compared to younger adults. This model received support from several fMRI and PET studies, demonstrating less lateralized or even balanced bilateral brain activity in older as compared to younger adults in WM tasks with both verbal and spatial stimuli (e.g., Cabeza, Anderson, Locantore, & McIntosh, 2002; Dixit, Gerton, Kohn, Meyer-Lindenberg, & Berman, 2000; Reuter-Lorenz et al., 2000). These reductions in lateralization with age were attributed to additional recruitment of the non-dominant hemisphere to compensate for age-related deteriorations in the dominant hemisphere (Cabeza, 2002). Alternatively, others suggested that reduced hemispheric lateralization reflects non-selective aging-induced reductions in the specificity of processing pathways (e.g., dedifferentiation; see Li & Lindenberger, 1999; Li & Sikström, 2002).

Over the last decade, many electrophysiological studies of visual WM (VWM) used paradigms that presented visual stimuli bilaterally in both hemifields, but made use of the fact that the contralateral organization of the visual system leads to input from each hemifield being processed preferentially in the contralateral hemisphere. When participants memorize stimuli in one hemifield while information is presented bilaterally (i.e., the perceptual input in the two hemispheres is the same), differences in activity between the contra- and ipsilateral hemisphere can be attributed to memory-related processes (Luck & Vogel, 2013; Vogel & Machizawa, 2004). The difference between contra- and ipsilateral activity during the delay period of VWM tasks, the so-called contralateral delay activity (CDA, Vogel & Machizawa, 2004)^1^, was shown to vary with the number of items to be maintained in VWM (Luria, Balaban, Awh, & Vogel, 2016). This negative-going difference wave typically starts about 200 ms to 300 ms after the onset of the memory array, with the highest amplitudes at posterior parietal and lateral occipital electrode sites (Vogel & Machizawa, 2004). The CDA was originally described and studied in change-detection tasks using visuo-spatial configurations of colored squares or orientation bars (see Luck & Vogel, 1997; Vogel & Machizawa, 2004;). However, the CDA has also been observed in response to letters presented as memoranda in change-detection (Raisic, Burton, & Woodman, 2019) and verbal recall VWM tasks (Wiegand, Töllner, Habekost, et al., 2014). In line with a decline of VWM performance in older adults, CDA amplitudes have been shown to be reduced or less modulated by load in older compared to younger adults in both paradigms (Jost et al., 2011; Sander, Werkle-Bergner, & Lindenberger, 2011; Wiegand, Töllner, Dyrholm, et al., 2014; Wiegand et al., 2018).

The reader should note that in these paradigms, VWM performance and the CDA were typically calculated based on averaging across left and right hemifield conditions, thus disregarding any differences in hemispheric processing. Only very few studies have considered hemisphere-specific effects in the CDA, and their results were inconclusive. In a visuo-spatial change detection paradigm, Machizawa, Goh, Driver, and Husain (2012) found that the CDA decayed faster on right-hemifield trials (with processing occurring in the left hemisphere) than on left-hemifield trials (with processing in the right hemisphere). By contrast, McCollough et al. (2007) found a slightly, but not significantly, higher CDA amplitude in the right hemisphere compared to the left hemisphere. To our knowledge, no study has investigated hemispheric differences in the CDA in a VWM task that uses verbal stimuli, as yet. However, behaviorally (measured in terms of the capacity parameter K; e.g., Bundesen, 1990), a right-hemifield advantage was reported in both younger (Kraft et al., 2015) and older adults (Wiegand et al., 2018), and this lateralization was shown to be increased by right frontal and parietal transcranial direct current stimulation (tDCS) in older adults (Brosnan et al., 2017).

In the present study, we investigated age differences in material-dependent hemispheric lateralization in behavioral (K parameter; e.g., Bundesen, 1990; Cowan, 2001) and neural measures of VWM capacity (CDA), respectively. We re-analyzed data from two previously published studies that examined age differences in a spatial (Sander et al., 2011) and a verbal VWM task (Napiórkowski et al., 2019). According to theories of functional lateralization (Smith & Jonides, 1998), we expected a material-dependent hemispheric specialization in younger adults, with higher K estimates and larger CDA amplitudes for spatial stimuli presented in the left hemifield (i.e., processed in right hemisphere) compared to the right hemifield, and vice versa for verbal stimuli, that is, higher K estimates and larger CDA amplitudes for stimuli presented in the right hemifield (i.e., processed in the left hemisphere) than for those presented in the left hemifield. In line with the HAROLD model (Cabeza, 2002), we expected this material-dependent hemispheric lateralization in VWM to be reduced in older adults compared to younger adults.

## Methods

### Samples

Two previously published data sets were analyzed. The first sample consisted of 31 healthy younger adults (YA: *M*_age_ = 26.00, range 20–33 years; 18 female/13 male) and 29 healthy older adults (OA: *M*_age_ = 71.75, range 65–78 years; 11 female/18 male), both with data collected at Ludwig-Maximilians-Universität München (LMU Munich). Participants’ VWM was tested using a lateralized whole report task with letters as stimuli (see Duncan, 1999), henceforth referred to as the *verbal* VWM task. In addition, all participants were tested for crystallized intelligence using the Mehrfachwahl-Wortschatz-Test B (MWT-B; Lehrl, Merz, & Burkhard, 1977) and processing speed using the Trail Making Test A (Rodewald et al., 2012). Furthermore, older participants were tested for cognitive deficits with the Mini Mental State Examination (MMSE; Folstein, Folstein, & McHugh, 1975) and were only included if their score was above 26. Table 1 shows the cognitive (MWT-B and Trail Making Test) measures for a comprehensive sample description.

**Table 1.** Cognitive measures of the LMU sample performing the verbal task.

| Cognitive measures | Younger adults ($n = 31$) | Older adults ($n = 29$) |
| --- | --- | --- |
| | $M (SD)$ | $M (SD)$ |
| MWT-B | 29.71 (4.38) | 33.57 (1.60) |
| Trail Making Test A (in seconds) | 22.23 (8.30) | 43.92 (9.53) |

*Note.* $M$ = mean; $SD$ = standard deviation

The second sample consisted of 12 healthy younger adults (YA: *M*_age_ = 23.6, range 20– 25 years; 6 female/6 male) and 22 healthy older adults (OA: *M*_age_ = 72.8, range 70–75 years; 11 female/11 male), with data collected at the Max Planck Institute for Human Development (MPIB) in Berlin. Participants’ VWM was tested using a cued change detection task with colored squares at unique spatial positions, henceforth referred to as the *spatial* VWM task. Cognitive measures (upper half of Table 2) again included the MWT-B (Lehrl et al., 1977) as a test of crystallized intelligence and a test of processing speed, the Digit Symbol Substitution test (Wechsler, 1955). Furthermore, visual acuity was measured in Snellen decimal units using Landolt rings at 30 cm and 5 m distance (Geigy, 1977) (see lower half of Table 2).

**Table 2.** Cognitive measures of the MPIB sample performing the spatial VWM task.

| Cognitive measures | Younger adults ($n = 12$) | Older adults ($n = 22$) |
| --- | --- | --- |
| | $M (SD)$ | $M (SD)$ |
| MWT-B | 24.17 (3.69) | 29.23 (3.13) |
| Digit Symbol Substitution | 66.92 (10.67) | 48.77 (9.81) |
| Near vision | 0.82 (0.12) | 0.45 (0.15) |
| Far vision | 1.4 (0.55) | 1.02 (0.38) |

*Note.* $M$ = mean; $SD$ = standard deviation

Comparisons of the cognitive measure that was assessed in both samples, the MWT-B test for crystallized intelligence, revealed that in both age groups, the LMU participants had significantly higher values than the MPIB participants (YA: *t*(41) = –3.87, *p* < .001; OA: *t*(48) = –6.55, *p* < .001). Both data sets were collected in accordance with the Declaration of Helsinki on ethical principles, and all participants gave written informed consent prior to their participation.

### Tasks and Stimuli

Participants of the first sample performed a verbal VWM task (see also Wiegand et al., 2018), based on Bundesen’s (1990) computational Theory of Visual Attention (TVA). More specifically, they were asked to encode and retain (for later report) as many briefly presented arrays of letters while their electrophysiological and behavioral responses were recorded. The experimental paradigm is illustrated in Figure 1a.

**Figure 1.**
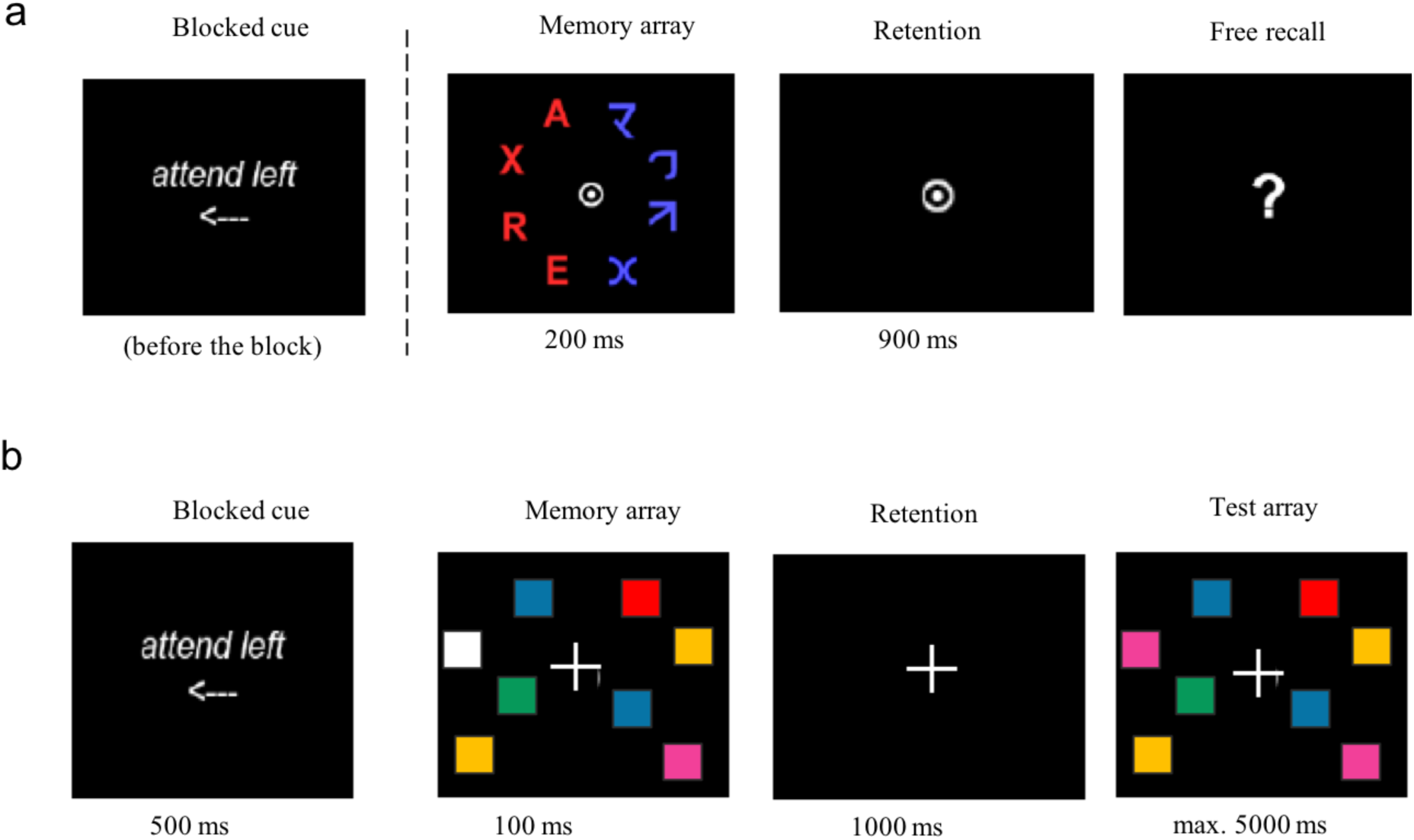
Verbal (a) and spatial (b) visual working memory task.

A cue was presented at the beginning of a block, prompting participants to pay attention to either the left or the right hemifield, while fixating a dot in the center of the screen. The cued hemifield was kept constant for a block of 40 consecutive trials, to avoid additional memory load. In each trial, a memory array consisted of four randomly chosen letters in the cued hemifield, four scrambled letters in the opposite hemifield, and a fixation dot in the middle. Participants had to encode and retain only the letters in the cued hemifield. The memory array was followed by a retention interval of 900 ms. Finally, a question mark appeared on the screen and participants had to verbally report as many letters as possible. The order and speed of reporting was irrelevant. A total of 220 trials, in which memory arrays were presented for 200 ms, were used for the CDA analyses. The experiment further contained 180 trials with individually adapted, shorter exposure durations. These trials were important for the TVA-based estimation of the attentional (performance) parameters, which was based on all 400 trials (Bundesen, 1990; Dyrholm et al., 2011; Kyllingsbæk, 2006; see also Napiórkowski et al., 2019, and Wiegand et al., 2018, for details of the EEG design in this task). Trial types were presented randomly intermixed. Performance feedback was given after each block.

Participants of the second sample performed a spatial VWM task (Luck & Vogel, 1997; Vogel & Machizawa, 2004): they were asked to retain arrays of colored squares presented on a display screen while their electrophysiological and behavioral responses were recorded. The experimental paradigm is illustrated in Figure 1b.

Each trial started with a visual cue presented centrally for 500 ms, telling participants to attend to either the left or the right hemifield while fixating a cross in the screen center. The cued hemifield was kept constant for 30 consecutive trials to avoid additional memory load. Next, a memory array consisting of a spatial configuration of four colored squares in each hemifield (on each side of the central fixation cross) was presented for 100 ms. Participants had to encode and retain the colored squares only in the cued hemifield. The memory array was followed by a retention interval of 1000 ms, after which a test array was presented for 5000 ms, which was either exactly identical to the memory array or differed in the color of one square in the cued hemifield. Participants had to indicate if the array had changed by pressing one of two buttons (change/no change). A total of 120 trials were presented. Note that participants completed other memory load and retention interval conditions as well, which are not analyzed in the present study as no matching conditions were available for the verbal VWM task (for more information, see Sander et al., 2011).

### Stimulus Presentation and Recordings

In the verbal VWM task, stimulus presentation was controlled with Eprime software (Psychology Software Tools Inc.). Behavioral responses were recorded manually by the experimenter, who also started the next trial via key press. Stimuli consisted of four red target letters (CIE values: 0.600, 0.327, 9.510) that were randomly chosen from the set [A, B, D, E, F, G, H, J, K, L, M, N, O, P, R, S, T, V, X, Z] and four equiluminant blue scrambled letters (CIE values: 0.190, 0.143, 9.660). Letters and scrambled letters appeared on an imaginary semicircle with a radius of 5.27° of visual angle on either the right or the left side of the fixation dot (size of the dot: 0.9° of visual angle in diameter). Diameters of letters and scrambled letters were 1.3° of visual angle. The same letter and symbol appeared only once in a given trial display. In five out of seven conditions, the letter array was followed by a (post-display) mask of eight red-blue scattered squares (size: 1.3° of visual angle) presented at each stimulus location (to curtail the effective stimulus exposure duration; together with the unmasked trials, this permitted the TVA-based fitting of the correct letter report over effective exposure duration function, from which the TVA attention parameters are estimated). Participants viewed the screen from a distance of 65 cm.

EEG data was recorded continuously with BrainAmp DC amplifiers (BrainVision Products GmbH) from 64 Ag/Ag-Cl electrodes, including two electrodes for horizontal EOG at the outer canthi and one electrode for vertical EOG. During recording, all electrodes were referenced to the FC_z_ electrode. Electrode impedances were maintained below 5 kΩ before recordings. The EEG was recorded with a band-pass filter of 0.1–250 Hz and digitized with a sampling rate of 1000 Hz (see also Napiórkowski et al., 2019).

In the spatial VWM task, stimulus presentation and recording of behavioral responses was controlled with Eprime v1.2 software (Psychology Software Tools Inc.). Stimuli consisted of colored squares (0.65° × 0.65° of visual angle) presented on a grey background (RGB values: 200, 200, 200) within an area of 4° × 7.3° of visual angle to the right and left of the central fixation cross (distance to the cross was 1.5°). The spatial locations of the squares were determined at random, with a minimum distance of 2° between the centers of adjacent squares. Colors were randomly selected from a set of 11 highly discriminable values: black (RGB values: 0, 0, 0), white (RGB: 255, 255, 255), grey (RGB: 126, 123, 126), blue (RGB: 0, 0, 255), green (RGB values: 0, 255, 0), red (RGB: 255, 0, 0), cyan (RGB: 0, 255, 255), violet (RGB: 255, 0, 255), brown (RGB: 153, 102, 51), orange (RGB: 255, 112, 1), and yellow (RGB: 255, 255, 0). The same color was not repeated more than twice per array. Eye to monitor distance was 70 cm.

EEG data was recorded continuously with BrainAmp amplifiers (BrainVision Products GmbH) from 64 Ag/Ag-Cl electrodes, including two electrodes for horizontal EOG at the outer canthi and one electrode for vertical EOG. During recording, all electrodes were referenced to the right mastoid electrode. Electrode impedances were maintained below 5 kΩ before recordings. The EEG was recorded with a band-pass filter of 0.1–250 Hz and digitized with a sampling rate of 1000 Hz (see also Sander et al., 2011).

### Analysis of Behavioral Data

For the analysis of behavioral responses, we computed one VWM capacity limit parameter, parameter K, per participant and hemifield. The computation of K differed between the two VWM tasks because of the different response formats (i.e., free recall in the verbal VWM task versus change/no-change decision indicated by button press in the spatial VWM task). Note, however, that the estimates of K in both tasks can be interpreted as the maximum number of objects maintained in VWM at a time.

Behavioral performance in the verbal VWM task was analyzed using a standard TVA-based fitting procedure (Bundesen, 1990; Dyrholm et al., 2011; Kyllingsbæk, 2006). In a maximum-likelihood procedure, individual’s response accuracy is modeled as a function of effective exposure duration. The function is described by four parameters: 1) the VWM capacity parameter K_verbal_, which is given by the asymptote of the function; 2) the processing speed parameter C, given by the slope of the function; 3) the perceptual threshold parameter t_0_, which is given by the x-axis intercept of the function; and 4) parameter µ, which indicates the persistence duration of iconic memory in unmasked display conditions (Wiegand, Töllner, Habekost, et al., 2014). Only parameter K_verbal_ was relevant for statistical analyses at issue in the present study. The quality of the fitting, calculated as the shared variance between the estimated and empirically obtained mean scores of correctly reported letters across all conditions, was good (*R*^*2*^ > 0.87 for all participants). We further calculated the correlation between K_verbal_ and the empirically obtained mean score in the 200 ms unmasked condition (i.e., the exposure time condition used for the CDA analysis), which also showed a high correspondence of *r* = 0.94.

In the spatial VWM task, we determined K_spatial_ using Cowan’s (2001) formula: (hit rate – false alarm rate) * set size. Hit rate was defined as the proportion of change trials on which a change was correctly reported. False alarm rate was defined as the proportion of no-change trials on which a change was incorrectly reported. Set size denotes the number of objects that were presented in the cued hemifield, which equaled 4 in the data analyzed in the present study.

### Analysis of Electrophysiological Data

For preprocessing, the EEG data of both samples were re-referenced to mathematically linked mastoids, downsampled to 250 Hz, and band-pass filtered between 0.5 and 100 Hz. Afterwards, data epochs were extracted from –2 s to 2 s with respect to the onset of the memory array. An independent component analysis (ICA) was used to correct for eye movement, heart beat artefacts, muscle activity, and noise. Independent components (ICs) representing artefactual sources were visually identified and removed from the data. Afterwards the data was visually inspected and trials still containing eye movements and muscle activity were excluded from the analysis.

For the verbal VWM task, up to four letters could be reported in the free recall. As partially correct trials were also considered valuable, all artefact-free trials were analyzed. The mean percentage of artifact-free trials was between 95.7 % (left hemifield) and 94.5 % (right hemifield) for younger adults, and between 90.0 % (left hemifield) and 90.6 % (right hemifield) for older adults. Data was baseline-corrected from –0.5 s to the onset of the memory array. This was possible because the hemifield cue was indicated at the beginning of each block and not at the beginning of each trial.

For the spatial VWM data, only trials with a correct change/no-change response were analyzed. The mean percentage of correct, artefact-free trials was between 70.5 % (left hemifield) and 65.5 % (right hemifield) for younger adults, and between 68.0 % (left hemifield) and 66.3 % (right hemifield) for older adults. Data was baseline-corrected from –1 s to –0.5 s. Baseline corrections needed to be done in this time window (i.e., not directly before the onset of the memory array), because the visual cue indicating the to-be-attended hemifield appeared on the screen at –0.5 s.

We calculated the CDA amplitude as the difference between the activity contralateral and ipsilateral to the attended hemifield for each subject. Visual inspection of the topography of the CDA in the verbal and spatial VWM task showed strongest activation at posterior and occipital electrodes (Figure 2). For our subsequent analyses, to maximize the probability to detect hemispheric differences in lateralization, we focused on the electrode that showed the strongest CDA amplitudes in each task and sample (EOI). In line with previous research (e.g., Luria, 2016), this was the electrode PO_7/8_.

**Figure 2.**
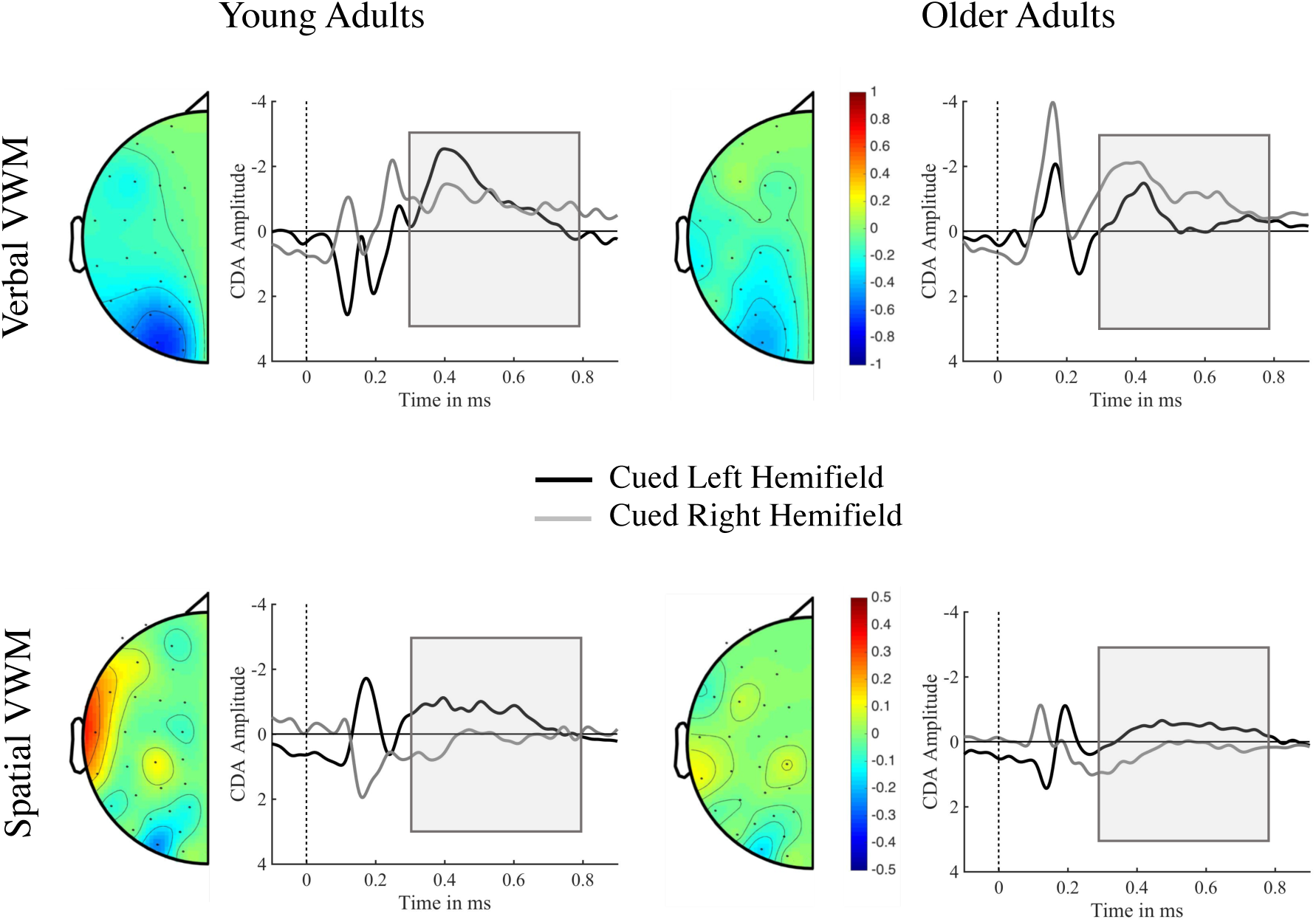
Contralateral delay activity (CDA). Topography of hemispheric lateralization effects, independent of presentation hemifield, and grand average event-related potentials (ERPs) at PO_7/8_ for left- and right-cued trials, separately for younger adults (left panel) and older adults (right panel). In both tasks and age groups, maximum hemispheric lateralization is observed over posterior electrodes, i.e., PO_7/8._ The task-specific time windows of interest are highlighted in the ERP curves. Upper panels: Verbal visual working memory (VWM) task. Data was baseline-corrected from –0.5 s to 0 s. Lower panels: Spatial VWM task. Data was baseline-corrected from –1 s to –0.5 s.

As presentation times of memory arrays differed between tasks, and as previous studies point to possible differences between the spatial and verbal tasks in the timing of the maximal CDA amplitudes (Sander et al., 2011; Wiegand et al., 2014, 2018), we tested each sample point within a broader time window of 300 ms to 800 ms for a reliable difference between ipsi-and contralateral activity (collapsed across hemispheres and age groups, but separately for each task), using a correction procedure for multiple comparisons based on false discovery rates (Benjamini-Yekutieli procedure, as implemented in FieldTrip). This procedure yielded significant differences between the ipsi- and contralateral activity between 320 ms and 660 ms for the verbal VWM task and between 410 ms and 750 ms for the spatial VWM task. Figure 2 shows the task-specific event-related potential curves.

### Statistical Analysis

To account for the unbalanced design, we used the lme4 package (Bates, Maechler, Bolker, and Walker, 2015) in R (R Core Team, 2018) to perform linear mixed effects analysis of the effects of task, age group, and hemifield on the two dependent variables, namely K parameter and CDA mean amplitude. As random effects, we had intercepts for participants. As fixed effects, we tested the effects of task (verbal/spatial), age group (younger/older adults) and to-be-attended hemifield (left/right) separately against a null model including only a random intercept for each participant. To test whether hemispheric lateralization was modulated by task, we also tested a model including the interaction against the full main effect model. Finally, to test whether the hemispheric lateralization by task interaction differed by age group, we tested the model with the three-way interaction against the model with the two-way interaction. P-values were obtained by likelihood ratio tests of the model with the effect in question against the model without the effect in question. Effects with *p* < .05 were considered significant. Significant effects were followed up by post-hoc comparisons using t-tests or Wilcoxon tests when the assumption of normality was violated.

## Results

### Behavioral Results

The linear mixed effects model on the behavioral level revealed a main effect of age group (χ^2^ (1) = 11.32, *p* = 0.00) with overall better performance for younger adults (K_YA_: *M* = 3.06, *SD* = 0.63) compared to older adults (K_OA_: *M* = 2.60, *SD* = 0.70), and a main effect of task (χ^2^ (1) = 56.26, *p* = 0.00) with overall better performance for the verbal task (K_verbal_: *M* = 3.15, *SD* = 0.45) compared to the spatial task (K_spatial_: *M* = 2.20, *SD* = 0.67). The main effect of hemifield was not significant (χ^2^ (1) = 1.83, *p* = 0.18), but we found a reliable interaction of hemifield and task (χ^2^ (1) = 17.15, *p* = 0.00). In line with our hypothesis of a hemispheric lateralization for verbal and spatial material, we observed a higher K for left-hemifield compared to right-hemifield stimuli in the spatial task (K_left_: *M* = 2.28, *SD* = 0.72, K_right_: *M* = 2.13, *SD* = 0.62), and a higher K for right-hemifield stimuli compared to left-hemifield stimuli in the verbal task (K_left_: *M* = 3.07, *SD* = 0.46, K_right_: *M* = 3.24, *SD* = 0.42).

Finally, we observed that age group modulated the two-way interaction between hemifield and task as indicated by a significant three-way interaction (χ^2^ (3) = 8.72, *p* = 0.03). We followed up on this interaction testing the effect of hemifield (i.e. the lateralization) within task and age groups separately: For younger adults, we found a reliable effect of lateralization in the spatial task (paired t-test, *t*(11) = 2.50, *p* = 0.03) and in the verbal task (Wilcoxon signed rank test, *V* = 128, *p* = 0.02). For older adults, we observed a reliable lateralization only in the verbal task (*t*(28) = –5.39, *p* = 0.00), but not in the spatial task (*t*(21) = 0.59, *p* = 0.56). See Figure 3 for a summary of the results.

**Figure 3.**
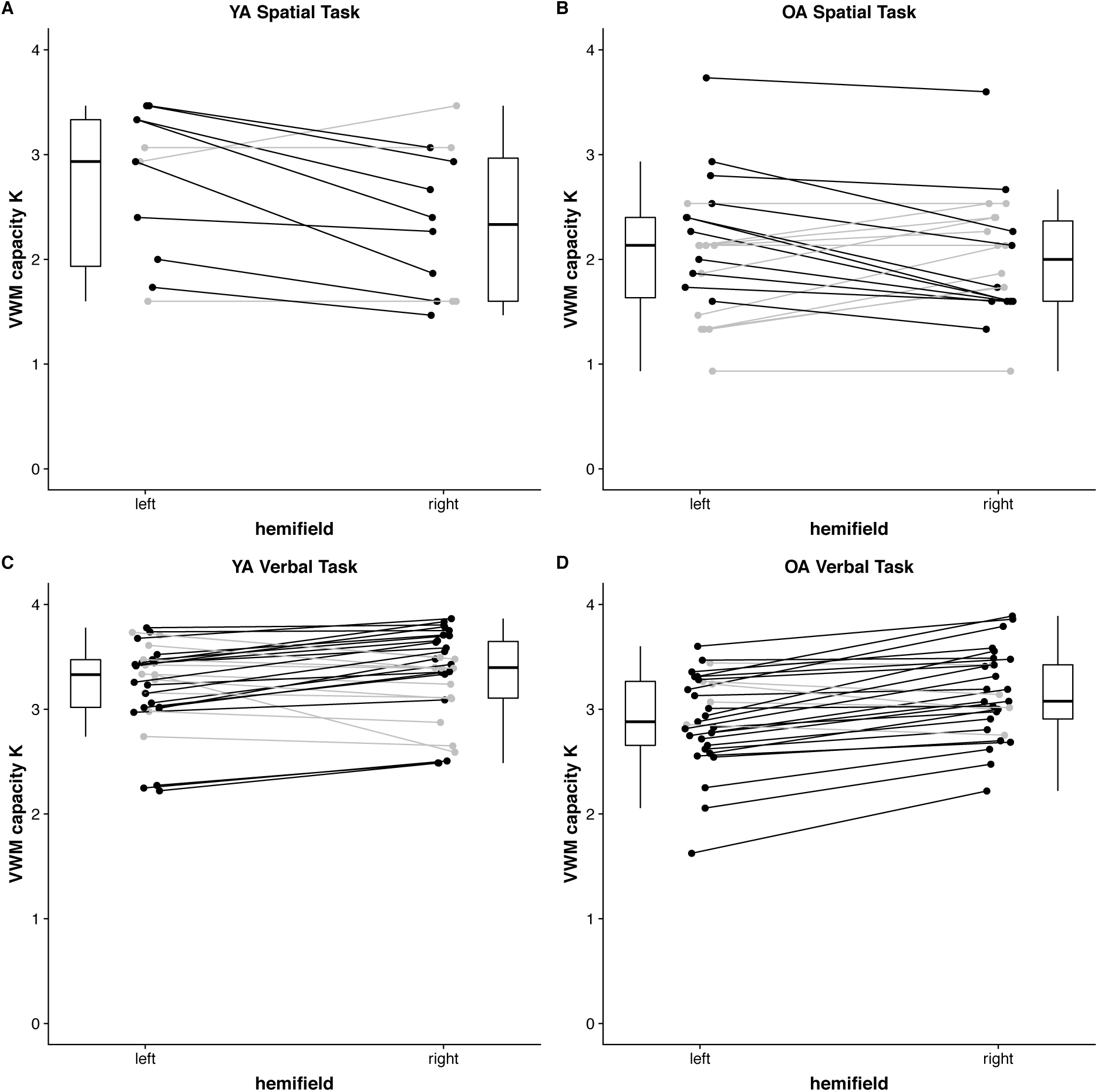
Summary of behavioral results. A & B: Spatial task. Both younger adults (A) and older adults (B) show higher VWM capacity (indicated by K values) for left-hemifield stimuli compared to right-hemifield stimuli. C & D: Verbal task. Both younger adults (C) and older adults (D) show higher visual working memory (VWM) capacity (indicated by K values) for right-hemifield stimuli compared to left-hemifield stimuli. Individual participants are represented by dots, paired measures are connected by lines. Participants displaying an effect in the expected direction are depicted in black, whereas those with opposite patterns or no difference are depicted in gray. Boxplots are used to indicate the distribution of the data, visualizing the median, first, and third quartiles (lower and upper hinges). YA: younger adults; OA: older adults.

### Electrophysiological Results

The linear mixed effects model on the neural level (i.e., CDA amplitudes) revealed a main effect of task (χ^2^(1) = 14.78, *p* = 0.00) with larger (thus, more negative-going) CDA amplitudes in the verbal task (CDA_verbal_: *M* = -.90, *SD* = 1.63) compared to the spatial task (CDA_spatial_: *M* = -.16, *SD* = 0.53) and a reliable two-way interaction of hemifield and task (χ^2^(1) = 6.23, *p* = 0.01). More specifically, CDA amplitudes were larger (thus, more negative-going) for left-hemifield stimuli than for right-hemifield stimuli in the spatial task (CDA_left_: *M* = -.46, *SD* = 0.44, CDA_right_: *M* = .25, *SD* = 0.34), but larger (thus, more negative-going) for right-hemifield stimuli than for left-hemifield stimuli in the verbal task (CDA_left_: *M* = -.75, *SD* = 1.54, CDA_right_: *M* = –1.05, *SD* = 1.72). Post-hoc comparisons revealed a reliable lateralization effect in the spatial task (Wilcoxon signed rank test, *V* = 10, *p* = 0.00), but not in the verbal task (Wilcoxon signed rank test, *V* = 961, *p* = 0.74). There was no main effect of age group nor an interaction involving age group (all *p* > .18). Figure 4 provides a summary of the results.

**Figure 4.**
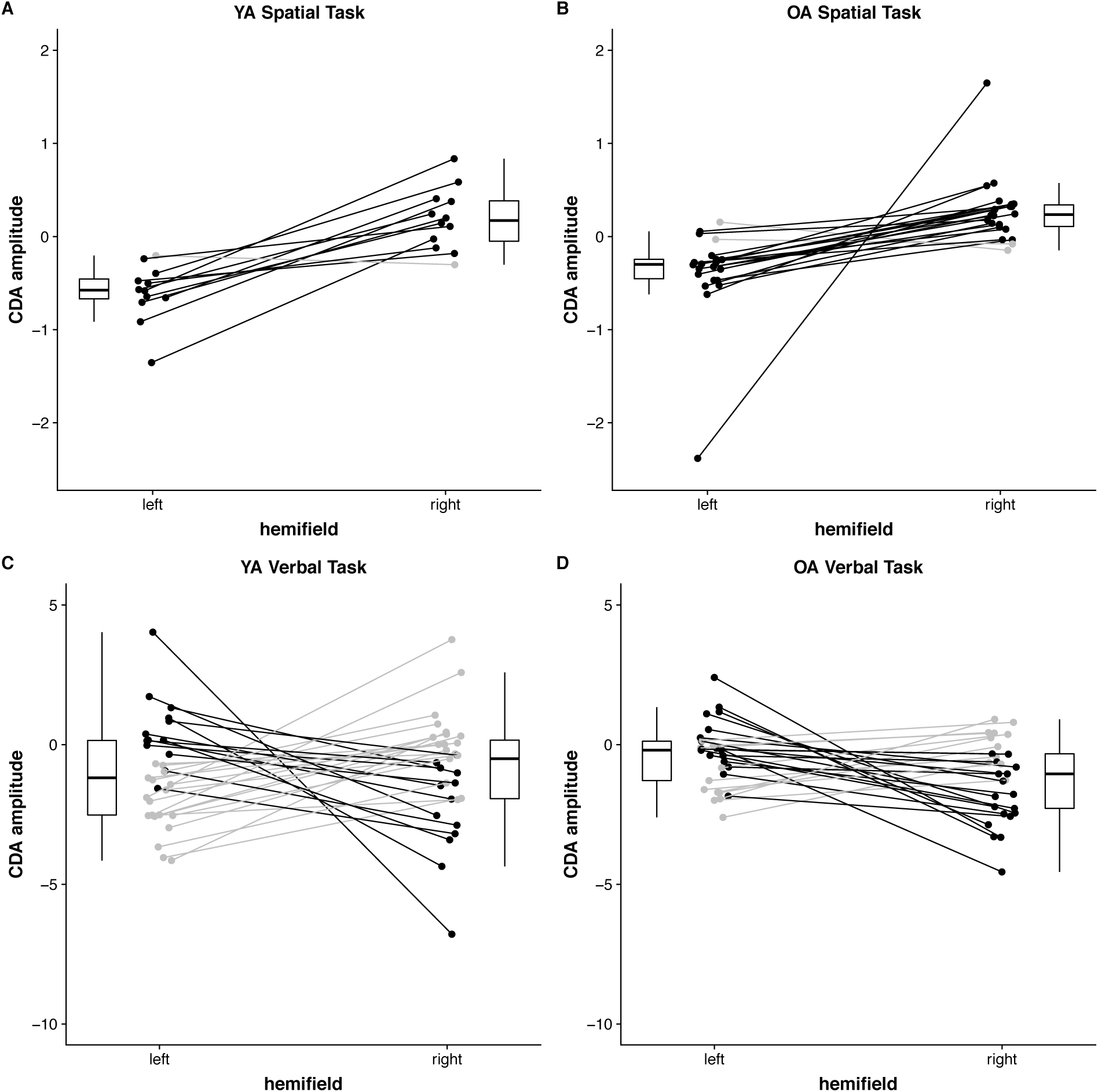
Summary of EEG results. A & B: Spatial task. Both younger adults (A) and older adults (B) show larger negative contralateral delay activity (CDA) amplitudes for left-hemifield stimuli than for right-hemifield stimuli. C & D: Verbal task. There is no clear lateralization pattern observable in either younger adults (C) nor older adults (D). Individual participants are represented by dots, paired measures are connected by lines. Participants displaying an effect in the expected direction are depicted in black, whereas those with opposite patterns or no difference are depicted in gray. Boxplots are used to indicate the distribution of the data, visualizing the median, first, and third quartiles (lower and upper hinges). YA: younger adults; OA: older adults.

## Discussion

In the present study, we investigated age differences in hemispheric lateralization effects on behavioral and neural markers of VWM. According to theories of functional lateralization (Smith & Jonides, 1998), we expected to find material-specific (spatial vs. verbal) hemispheric lateralization during VWM performance in younger adults. In line with the HAROLD model (Cabeza, 2002), and with notions of aging-induced losses in processing fidelity and specificity (e.g., Li & Sikström, 2002), we hypothesized that this hemispheric lateralization would be reduced in older adults.

On the behavioral level, our findings are in line with a right-hemispheric specialization for the maintenance of spatial information, and a left-hemispheric specialization for the maintenance of verbal information (Machizawa et al., 2012; Reuter-Lorenz et al., 2000; Smith & Jonides, 1998; Spotorno & Faure, 2011; Thomason et al., 2008). In the spatial task, performance was higher for stimuli presented in the left hemifield, whereas in the verbal task performance was higher for stimuli presented to the right hemifield. This task-dependent hemispheric lateralization differed between younger and older adults: Whereas younger adults showed a reliable lateralization with a preference in the direction of the expected hemisphere in both tasks, older adults showed a clear lateralization only in the verbal, but not in the spatial task.

In line with the behavioral results, we found evidence for hemispheric lateralization also on the neural level, as indicated by a reliable interaction of hemifield and task, although post-hoc tests revealed that the hemispheric lateralization was reliable in the spatial VWM task, but not in the verbal VWM task. In the spatial task, larger CDA amplitudes were observed for stimuli processed in the right hemisphere versus those processed in the left hemisphere. This finding is in accordance with previous reports of a right-hemispheric dominance for spatial VWM processes (Reuter-Lorenz et al., 2000; Smith & Jonides, 1998; Thomason et al., 2008) and extends the previous results obtained from PET and fMRI studies to the main *electrophysiological* marker of VWM capacity, the CDA. While previous neuroimaging studies also showed a clear left-hemispheric lateralization of the maintenance of verbal information (Reuter-Lorenz et al., 2000; Smith & Jonides, 1998; Thomason et al., 2008), the CDA in the verbal task did not differ for stimuli presented to the right versus left hemifield. The difference in hemispheric lateralization between the two tasks probably reflects differences in the type of maintenance processes that participants employ: While in the verbal task the reliance on verbal rehearsal strategies may induce hemispheric lateralization in rather prefrontal regions, maintenance of spatial memoranda may rely more on posterior parietal areas, in which the main generator sources of the CDA are assumed to be located (Becke, Müller, Vellage, Schoenfeld, Hopf, 2015; Eriksson et al., 2015; Luck & Vogel, 2013). Thus, the CDA may capture hemispheric lateralization more reliably in a task that relies heavily on posterior parietal regions.

Importantly, hemispheric lateralization on the neural level did not differ between age groups. Thus, our results argue against a general reduction of hemispheric lateralization with aging (Cabeza, 2002). It is possible that the age-related reduction in neural hemispheric asymmetries occurs primarily in prefrontal areas (Cabeza et al., 2002; Reuter-Lorenz et al., 2000), may not generalize to posterior parietal brain regions, and is therefore not captured by the CDA. In contrast to the CDA results, on the behavioral level, we observed hemispheric lateralization in older adults in the verbal task but not in the spatial task. Thus, our results support differential lateralization effects in VWM depending on the type of material that has to be maintained (Höller-Wallscheid, Thier, Pomper, & Linder, 2016) and indicate that those can vary with aging. This is in accordance with previous studies that reported a shallower age-related decline in verbal as compared to spatial WM tasks (Jenkins et al., 2000; Myerson et al., 2003). This finding would also be in line with the RHAM (Brown & Jaffe, 1975) suggesting that functions associated with the right hemisphere (i.e., spatial WM) show a faster age-related decline than functions associated with the left hemisphere (i.e., verbal WM). However, with the current study, it is not possible to differentiate between effects that are due to differences between domains and effects that are due to genuine neural differences between hemispheres. To disentangle these effects, future studies could probe older adults’ neural systems with regard to hemispheric lateralization using stimuli presented laterally, for which no genuine hemispheric preference exists. Hemispheric differences independent of material-specific lateralization would more clearly speak to this question.

In sum, while we do not find clear evidence for age differences in hemispheric lateralization, our results do not necessarily preclude that aging induces a loss in processing fidelity and specificity (e.g., Li & Sikström, 2002). In particular, the observation that older adults did not show a clear lateralization pattern in the spatial task indicates that information may be processed differently in older compared to younger adults with downstream consequences for later memory performance (see also Sommer et al., 2019 for similar evidence regarding episodic memory). Future work is therefore needed to clarify under which conditions processing specificity—hemispheric lateralization being one example— changes across the lifespan.

## Acknowledgments

Data was collected at the Department of Psychology at Ludwig-Maximilians-Universität München, Munich, Germany, and within the project, “Cognitive and Neuronal Dynamics of Memory across the Lifespan (ConMem)” at the Center for Lifespan Psychology, Max Planck Institute for Human Development, Berlin, Germany. The Max Planck Society financially supported the research. The research at Ludwig-Maximilians-Universität München was supported by the European Union’s Seventh Framework programme for research, technological development, and demonstration under the Marie Sklodowska-Curie Initial Training Network grant agreement no. 606901.

MCS was further supported by the MINERVA program of the Max Planck Society. IW was supported by the European Union’s Horizon 2020 research and innovation programme, Marie Sklodowska-Curie Actions, under grant 702483. We thank all our student assistants for their support in data collection. We are grateful to Julia Delius for editorial assistance.

## Footnotes

1 The CDA is sometimes also referred to as the contralateral negative slow wave (CNSW; Klaver, Talsma, Wijers, Heinze, & Mulder, 1999), sustained posterior contralateral negativity (SPCN; Brisson & Jolicœur, 2007), or contralateral search activity (CSA; Emrich, Al-Aidroos, Pratt, & Ferber, 2009). For the sake of simplicity, we will only use the term CDA.

